# A genetic hazard score to personalize prostate cancer screening, applied to population data

**DOI:** 10.1101/619718

**Authors:** Minh-Phuong Huynh-Le, Chun Chieh Fan, Roshan Karunamuni, Eleanor I. Walsh, Emma L. Turner, J. Athene Lane, Richard M. Martin, David E. Neal, Jenny L. Donovan, Freddie C. Hamdy, J. Kellogg Parsons, Rosalind A. Eeles, Douglas F. Easton, ZSofia Kote-Jarai, Ali Amin Al Olama, Sara Benlloch Garcia, Kenneth Muir, Henrik Gronberg, Fredrik Wiklund, Markus Aly, Johanna Schleutker, Csilla Sipeky, Teuvo LJ Tammela, Børge G. Nordestgaard, Tim J. Key, Ruth C. Travis, Paul Pharoah, Nora Pashayan, Kay-Tee Khaw, Stephen N. Thibodeau, Shannon K. McDonnell, Daniel J. Schaid, Christiane Maier, Walther Vogel, Manuel Luedeke, Kathleen Herkommer, Adam S. Kibel, Cezary Cybulski, Dominika Wokolorczyk, Wojciech Kluzniak, Lisa Cannon-Albright, Hermann Brenner, Ben Schöttker, Bernd Holleczek, Jong Y. Park, Thomas A. Sellers, Hui-Yi Lin, Chavdar Slavov, Radka Kaneva, Vanio Mitev, Jyotsna Batra, Judith A. Clements, Amanda Spurdle, Australian Prostate Cancer BioResource (APCB), Manuel R. Teixeira, Paula Paulo, Sofia Maia, Hardev Pandha, Agnieszka Michael, Andrzej Kierzek, Ian G. Mills, Ole A. Andreassen, Anders M. Dale, Tyler M. Seibert, The PRACTICAL Consortium

## Abstract

**Background:** Genetic risk stratification may inform decisions of whether—and when—a man should undergo prostate cancer (PCa) screening. We previously validated a polygenic hazard score (PHS), a weighted sum of 54 single-nucleotide polymorphism genotypes, for accurate prediction of age of onset of aggressive PCa and improved screening performance. We now assess the potential impact of PHS-informed screening.

**Methods:** United Kingdom population data were fit to a continuous model of age-specific PCa incidence. Using hazard ratios estimated from ProtecT trial data, age-specific incidence rates were calculated for percentiles of genetic risk. Incidence of higher-grade PCa (Gleason≥7) was estimated from age-specific data from the linked CAP trial. PHS and incidence data were combined to give a risk-equivalent age, when a man with a given PHS percentile will have risk of higher-grade PCa equivalent to that of a typical man at age 50 (50-years standard). Positive predictive value (PPV) of PSA testing was calculated using PHS-adjusted (PCa-risk-equivalent age) groups identified from ProtecT.

**Results:** Expected age of onset of higher-grade PCa is modulated by 19 years between the 1^st^ and 99^th^ PHS percentiles. A man with PHS in the 99^th^ percentile reaches 50-years-standard risk at age 41; conversely, a man in the 1^st^ percentile reaches this risk at age 60. PPV of PSA was higher for men with higher PHS-adjusted age.

**Conclusions:** PHS informs PCa screening strategies with individualized estimates of risk-equivalent age for higher-grade PCa. Screening initiation could be adjusted according to a man’s genetic hazard score, improving PPV of PSA screening.

## Introduction

Prostate cancer (PCa) is the second-most-common malignancy in men worldwide with nearly 1.3 million cases diagnosed globally in 2018 alone^1^. PCa was the third leading cause of European male cancer mortality in 2018, following mortality from lung and colorectal cancers^2^. PCa screening with prostate-specific antigen (PSA) testing can reduce mortality^3^, but universal screening may result in the overdetection of cancers that would never become clinically apparent in a man’s life-time and overtreatment of indolent disease. Guidelines recommend that individual men participate in informed decision making about screening, taking into account factors such as their age, race/ethnicity, family history, and preferences^4–6^.

Assessment of a man’s genetic risk of developing PCa has promise for guiding individualized screening decisions^7,8^. We previously developed a polygenic hazard score (PHS) as a weighted sum of 54 single-nucleotide polymorphism (SNP) genotypes using clinical and genetic data from 31,747 men^9^. Validation testing, performed in an independent dataset consisting of 6,411 men from the United Kingdom (UK) ProtecT study^10,11^, showed the PHS to be a significant predictor of age at diagnosis of aggressive PCa, defined as cases where any of the following applied: Gleason score ≥7, clinical stage T3-T4, PSA ≥10, or where there were nodal or distant metastases^9^. Risk stratification by the PHS also improved the screening performance of PSA testing; the positive predictive value of PSA testing for aggressive PCa increased as PHS increased^9^.

Here, we apply the PCa PHS to population data to assess its potential impact on individualized screening. Specifically, we combine genetic risk, measured by PHS, and known population incidence rates to estimate a risk-equivalent age: e.g. the age at which a man with a given PHS will have the same risk of higher-grade PCa as a typical man at age 50 years. Such genetic risk estimates can guide individualized decisions about whether—and at what age—a man might benefit from PCa screening.

## Methods

### Polygenic hazard score (PHS)

Full methodologic details of the development and validation of the PCa PHS have been described previously^9^. Briefly, the PHS was created using data from men of European ancestry as a continuous survival analysis model^12^ to predict the age of PCa onset. PHS was calculated as the vector product of a patient’s genotype (*X*_*i*_) for *n* selected SNPs and the corresponding parameter estimates (*β*_*i*_) from a Cox proportional hazards regression (equation 1):

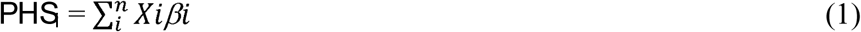

The 54 SNPs included in the model, and their parameter estimates, have been published^9^.

### Population age-specific incidence

Age-specific PCa incidence data were obtained for men aged 40-70 years from the United Kingdom, 2013-2015 (Cancer Research UK)^13^. Men may be less likely to receive PCa screening outside this age range^3,14^. The log of the PCa incidence data were fit using linear regression to develop a continuous model of age-specific PCa incidence in the UK (*h*_all_).

The UK age-specific proportion of incidence classified as higher-grade PCa was determined using data from the Cluster Randomized Trial of PSA Testing for Prostate Cancer (CAP) screening trial, which included men aged 50-69 at randomization^15^. Higher-grade PCa was defined for the present study as Gleason score ≥7. Men in the CAP trial were divided into 5-year age intervals at PCa detection, and the proportion of higher-grade disease in each age interval was calculated as the number of PCa diagnoses with Gleason score ≥7, divided by the total number of PCa diagnoses with known Gleason score in the CAP cohort^15^. The total (all ages) proportion of higher-grade PCa was also calculated from CAP data. The age-specific PCa curve, *h*_all_, was multiplied, within each 5-year age range, by the corresponding age-specific proportion of CAP diagnoses with Gleason score ≥7, to yield a continuous estimate of age-specific, *higher-grade* PCa incidence (*h*_higher-grade_). Of note, some men may have had Gleason score 6 with other clinically-significant features (i.e., high PSA, T3 stage, or regional/metastatic disease). Such men would also have been included in the definition of aggressive disease used in the PHS validation, despite having Gleason score <7, so the proportion used here is a conservative estimate of clinically-significant disease^9,16,17^.

### Impact of genetic risk on higher-grade PCa incidence

Men in ProtecT were categorized by PHS percentile ranges (0-2, 2-10, 10-30, 30-70, 70-90, 90-98, and 98-100) to correspond to percentiles of interest (1, 5, 20, 50, 80, 95, and 99, respectively). These percentiles refer to the distribution of PHS in the ProtecT dataset within controls aged <70. Hazard functions were calculated for each percentile range (*h*_percentile_) for aggressive PCa using Cox proportional hazards regression (parameter estimate, *ß*), following the methods published previously^9^. The reference for each hazard ratio (HR) was taken as the mean PHS among those men with approximately 50^th^ percentile for genetic risk (i.e., 30^th^-70^th^ percentile of PHS, called *PHS*_median_), and this *median* group was assumed to have a hazard for higher-grade disease matching the overall population (*h*_higher-grade_, calculated previously). Hazards for the other percentiles of interest (*h*_percentile_) were then calculated by determining the mean PHS among men in the corresponding percentile range (called *PHS*_percentile_) and applying equation 2:

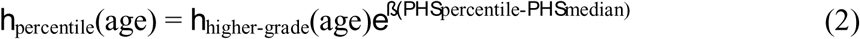

As described in the original validation of this PHS model for PCa^9^, PHS calculated in the ProtecT dataset will be biased by the disproportionately large number of cases included, relative to incidence in the general population. Leveraging the cohort design of the ProtecT study^10^, we therefore applied a correction for this bias, using previously published methods^18^ and the R ‘survival’ package (R version 3.2.2)^19,20^. The corrected PHS values were used to update *PHS*_percentile_ and *PHS*_median_ used in equation 2. Then, 95% confidence intervals for the HRs for each percentile were determined by bootstrapping 1,000 random samples from the ProtecT dataset, while maintaining the same number of cases and controls from the original dataset. The *h*_percentile,_ predicted partial hazard (product of *PHS*_percentile_ and the estimated ß), and standard errors (to account for sample weights) were calculated for each bootstrap sample.

Percentile-specific hazard estimates (*h*_percentile_) were visualized as the corresponding survival curves, where survival refers to survival free of higher-grade PCa diagnosis for men aged 50-70 years. Analogous HRs and survival curves were similarly calculated for the annualized incidence rates of *any* PCa, based on PHS for any PCa in the ProtecT dataset and the overall estimate of any PCa incidence in the UK (*h*_all_ instead of *h*_higher-grade_).

An individualized PHS to aid PCa screening decisions in the clinic might be facilitated by a readily interpretable translation of PHS to terms familiar to men and their physicians. The PHS was therefore combined with UK higher-grade PCa incidence data to give a risk-equivalent age, meaning when a man with a given PHS percentile would have the same risk of higher-grade PCa as, say, that of a typical man at 50 years old (50-years-standard risk). We defined ΔAge as the difference between age 50 and the age when PCa risk matches that of a typical 50-year-old man. 95% confidence intervals for the age when a man reaches 50-year-standard risk and ΔAge were determined using the HRs calculated from the 1,000 bootstrapped samples from ProtecT, described above.

Finally, we considered the common clinical scenario of a man presenting to his primary care physician (PCP) to discuss PCa screening. To illustrate how PHS might influence this discussion, we identified the subset of 945 men in the ProtecT validation dataset who were around the median age of 60 years (57-63), to represent a typical patient. From this subset, we created three groups: those whose PCa risk-equivalent age remained within the selected range (ages 57-63), those whose risk-equivalent age was <57, and those whose risk-equivalent age was >63. We then calculated the positive predictive value (PPV) of PSA testing for these three groups using methods described previously^9^; a true positive PSA was defined as biopsy-detected PCa (biopsy was indicated for men with PSA≥3.0 ng/mL). We calculated the PPV (and standard error [SE] of the mean) for PSA testing in the three PHS-adjusted (PCa risk-equivalent age) groups by taking 1,000 random samples of cases in the dataset, matching the ratio of controls to cases in ProtecT.

## Results

Linear regression yielded a model of PCa age-specific incidence rates (equation 5, *R*^2^=0.96 and *p*=0.001) that was highly consistent with empirical data reported by Cancer Research UK (**Figure 1**).

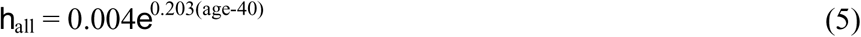

In the CAP study (of low-intensity PCa screening in the UK population)^15^, the proportion of PCa incidence classified as higher-grade disease was 59.7%, and the proportions of age-specific, high grade disease were: 34.3% for men aged 50-54, 45.6% for men aged 55-59, 52.5% for men aged 60-64, and 65.6% for men aged 65-69.

**Figure 1.**
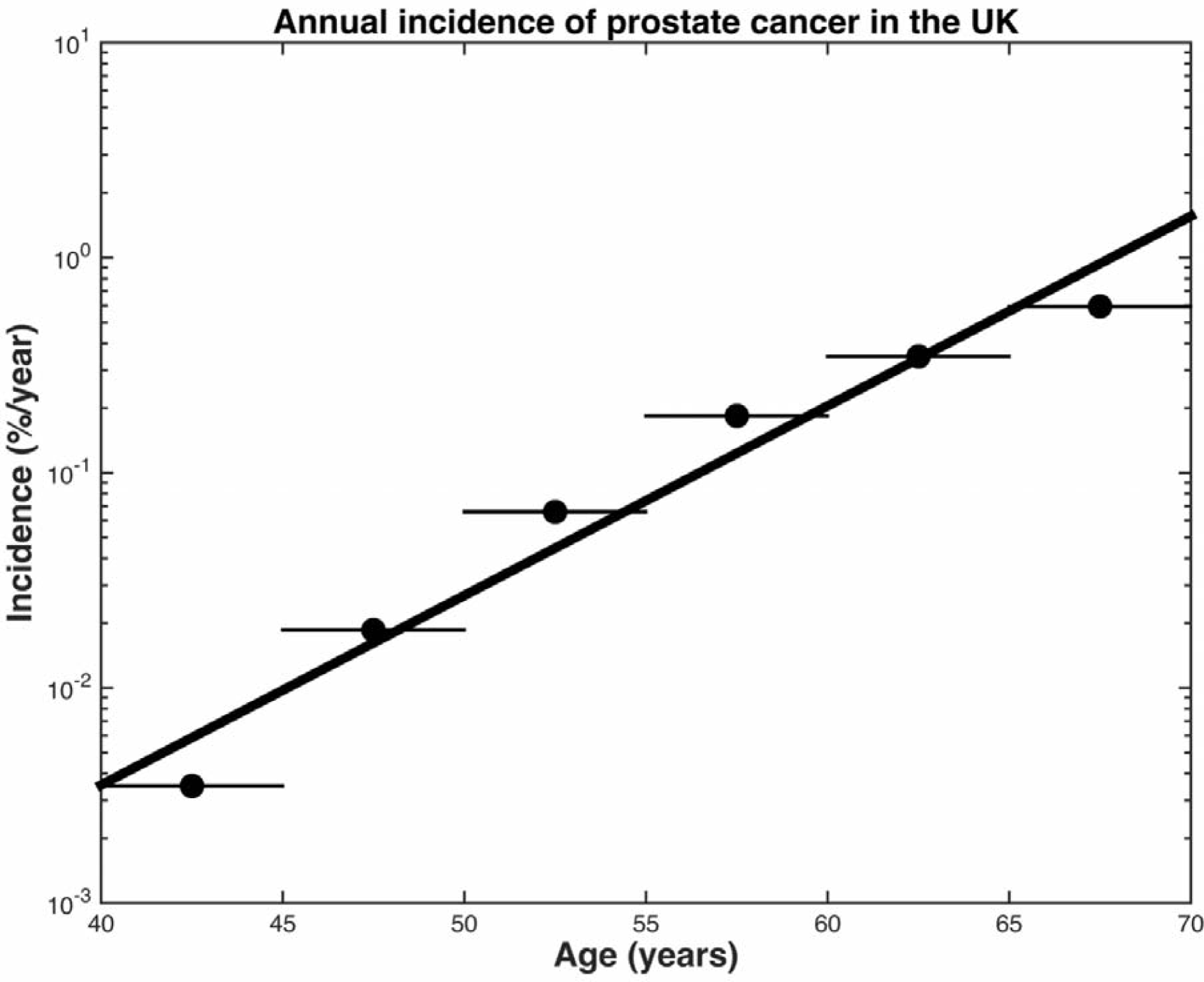
Annual incidence of prostate cancer in the United Kingdom, 2013-2015. Dots represent the raw, age-specific incidence rates of each age range, per 100,000 males. The black line represents the results of linear regression for an exponential curve to give a continuous model of age-specific incidence in the United Kingdom, *R*^2^=0.96, *p*=0.001.

Continuous estimates of survival free of higher-grade PCa are shown in **Figure 2** for various levels of genetic risk, as indicated by PHS percentile, showing a difference in age of onset related to PHS strata. **Supplemental eFigure 1** shows analogous results for survival free of *any* PCa. **Table 1** shows risk-equivalent age for each PHS percentile. The expected age of higher-grade PCa onset differs by 19 years between the 1^st^ and 99^th^ PHS percentiles. Specifically, a man with a PHS in the 99^th^ percentile reached a PCa detection risk equivalent to the 50-years standard at an age of 41 years. Conversely, a man with a PHS in the 1^st^ percentile would not reach the 50-years-standard risk level until age 60 years. Qualitatively, the curves for higher-grade PCa (**Figure 2)** and any PCa (**Supplemental eFigure 1**) maintain consistent horizontal shifts relative to curves for other PHS percentiles over the age range studied. Quantitatively, this was confirmed by ΔAge, which remained the same for each PHS percentile across a true age range of 40-70. Thus, ΔAge was taken to be approximately constant for each PHS percentile and is reported in **Table 1**.

**Table 1.**
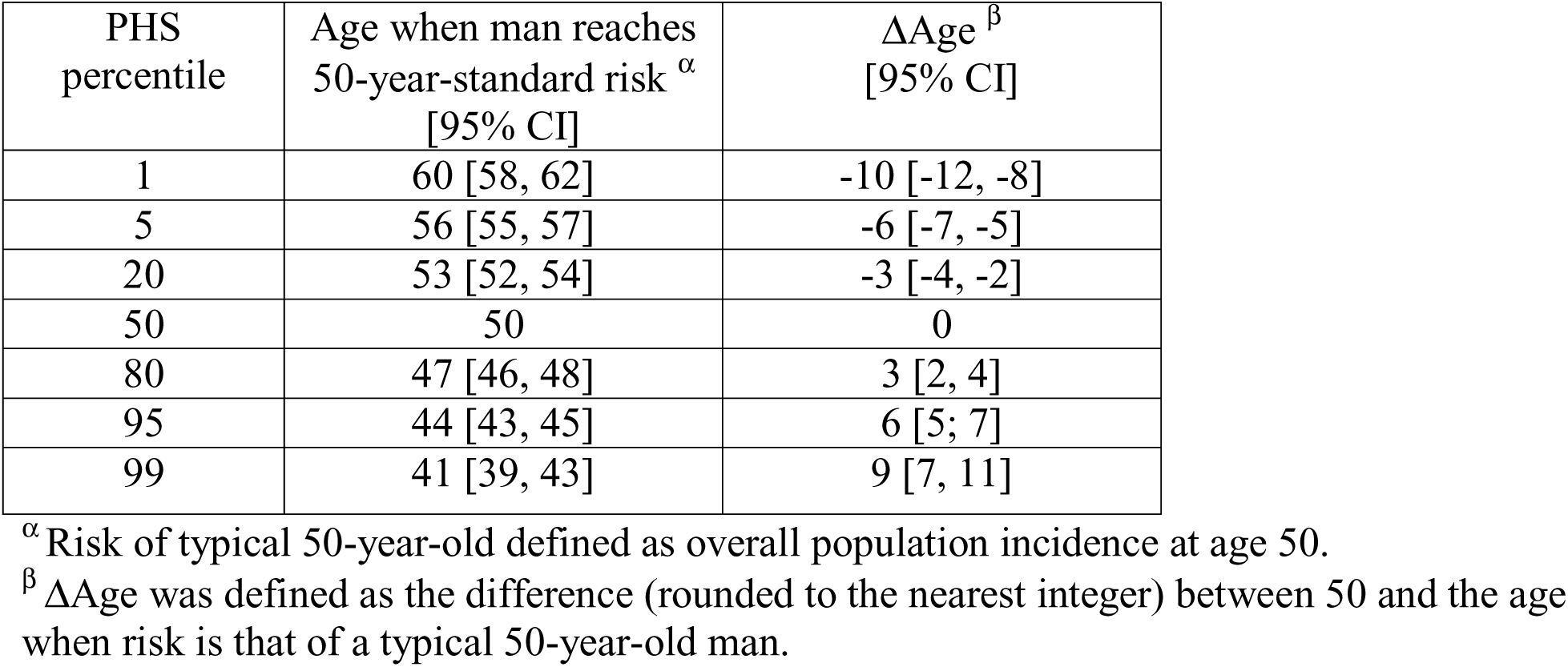
Risk-equivalent age for higher-grade prostate cancer, by polygenic hazard score (PHS) percentile.

| PHS percentile | Age when man reaches 50-year-standard risk <sup>α</sup><br>[95% CI] | ΔAge <sup>β</sup><br>[95% CI] |
| --- | --- | --- |
| 1 | 60 [58, 62] | -10 [-12, -8] |
| 5 | 56 [55, 57] | -6 [-7, -5] |
| 20 | 53 [52, 54] | -3 [-4, -2] |
| 50 | 50 | 0 |
| 80 | 47 [46, 48] | 3 [2, 4] |
| 95 | 44 [43, 45] | 6 [5, 7] |
| 99 | 41 [39, 43] | 9 [7, 11] |

<sup>α</sup> Risk of typical 50-year-old defined as overall population incidence at age 50.
<sup>β</sup> ΔAge was defined as the difference (rounded to the nearest integer) between 50 and the age when risk is that of a typical 50-year-old man.

**Figure 2.**
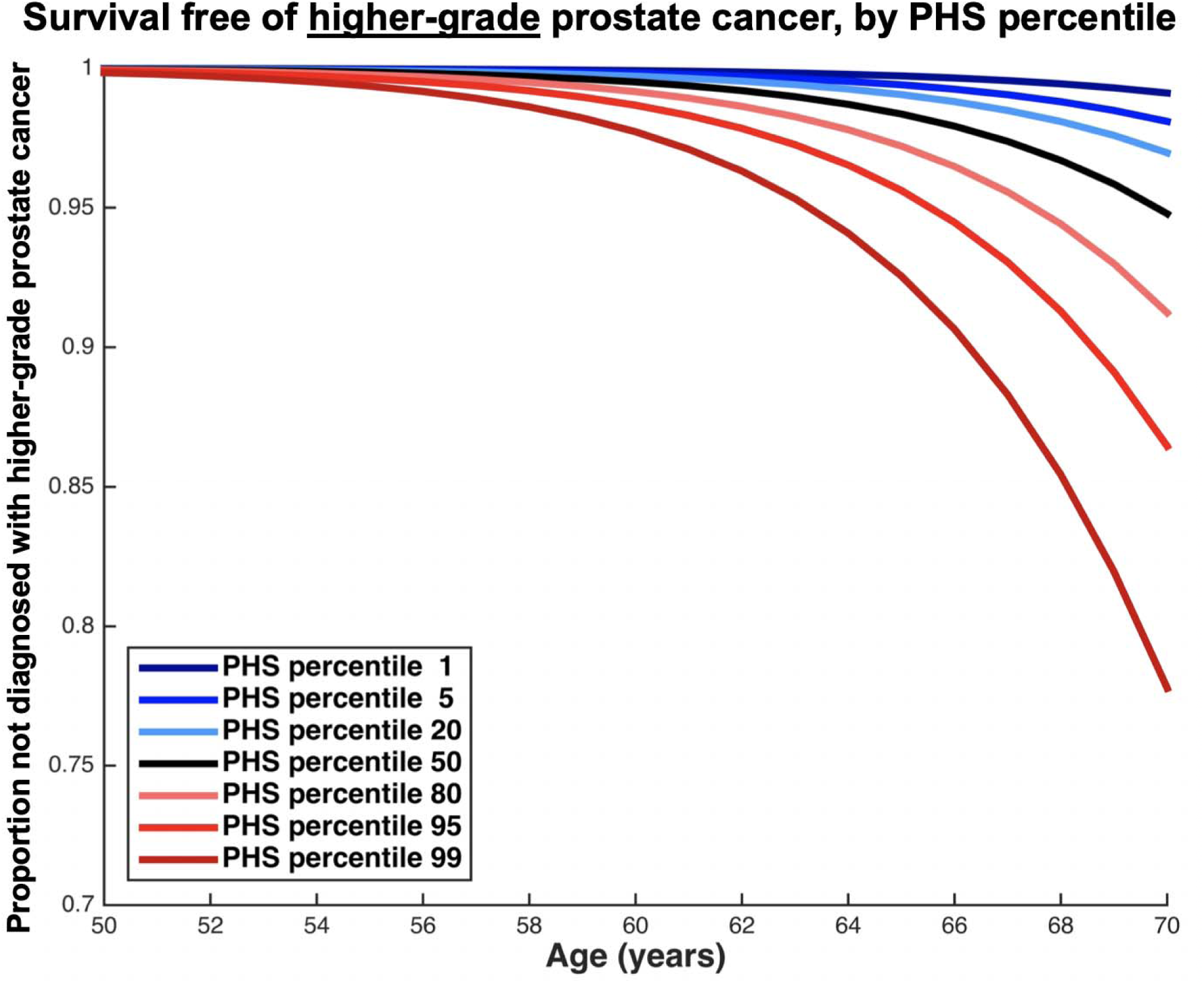
Survival free of higher-grade prostate cancer, as derived from application of polygenic hazard score (PHS) hazard ratios and population data from the United Kingdom. The overall population incidence is taken as the median risk (50^th^ percentile); this accounts for age-specific proportions of prostate cancer that was high grade in the CAP trial^15^. Hazard ratios were calculated within ProtecT data for various levels of genetic risk (i.e., PHS percentiles: 1, 5, 20, 80, 95, and 99) and used to adjust the median risk curve. Blue lines represent genetic risk lower than the median while red lines represent genetic risk higher than the median.

**Figure 3** shows the PPV of PSA testing (for any PCa) was 0.30 (SE: 0.009) for men approximately 60 years old (ProtecT men aged 57-63). PPV was lower for those with a PCa risk-equivalent age <57 years (0.10, SE: 0.05) and higher for those with PCa risk-equivalent age >63 years (0.48, SE: 0.03).

**Figure 3.**
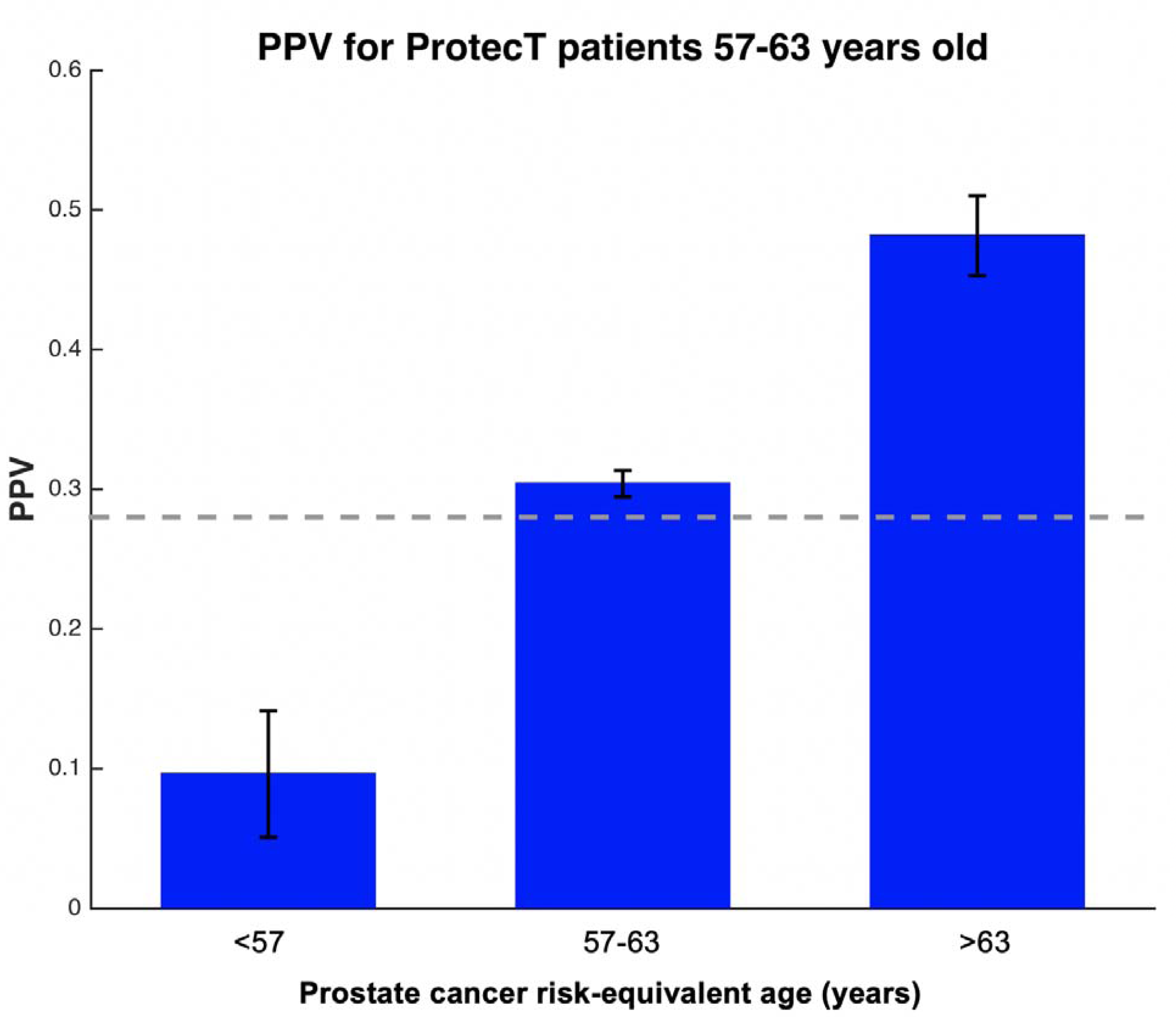
Application of prostate cancer risk-equivalent age to the clinical scenario of whether to screen a 60-year-old man (median age from ProtecT). The risk-equivalent age is the patient’s true age adjusted by PHS level. This plot shows results for all men from ProtecT aged approximately 60 years old (range: 57-63), grouped by their calculated prostate cancer risk-equivalent age: <57, 57-63, or >63. The positive predictive value (PPV) and the corresponding standard errors of the mean of PSA testing is shown for each of these 3 groups. The dashed, grey-line shows the expected PPV for PSA testing based on a published pooled analysis^5^.

## Discussion

We demonstrate the application of the PHS to population, age-specific incidence data to estimate age-specific risk of higher-grade PCa. The resulting age-specific hazard functions (displayed as survival curves in Figure 2) demonstrate clinically-meaningful differences across various levels of genetic risk (estimated by PHS). By combining these population curves with an individual’s genetic risk and true age, we demonstrate calculation of a risk-equivalent age for the onset of higher-grade PCa. This age relates a man’s current risk of PCa to that of the age-specific population average. The higher-grade PCa-free survival curves are modulated by 19 years between the 1^st^ and 99^th^ percentiles of PHS. Moreover, the PPV of PSA testing in three PHS-adjusted (PCa risk-equivalent age) groups demonstrated that PPV is significantly higher in men with higher risk-equivalent ages of onset of PCa. These results have important implications for clinicians considering discussions of whether—and when—to initiate PCa screening in an asymptomatic man.

It is well established that PCa can cause considerable mortality and morbidity, but if detected early, is a curable disease. PSA testing, while fast and convenient, remains an imperfect screening tool. PSA testing is associated with a small absolute decreased risk of death from PCa^3^, but carries a risk of overdetection and harm from overtreatment in men who would never have experienced clinical manifestations of their PCa^21^. Thus, universal screening for PCa comes at a high cost—both in burden on healthcare systems and in the sequelae arising from elevated PSA in men with indolent disease: unnecessary biopsy procedures, overdetection in men who may never have become symptomatic from PCa and treatment-related morbidities^4,5^. Conversely, there are some men who will develop higher-grade PCa and would benefit from screening, possibly even at a relatively young age. Screening guidelines recommend individualized decision-making, but the available quantitative or objective data to guide these decisions are insufficient. For instance, family history provides some guidance, but, genetic risk has been shown to be a more accurate predictor of age of aggressive PCa onset than patient-provided family history^9,22^. PHS, in conjunction with other informative factors such as family history, may help identify men who may develop those cancers that can cause morbidity and mortality^11^.

The stratification of men based on their genetic risk is of particular interest in the primary care setting, where the majority of PCa screening discussions take place. Shared decision-making between patient and physician has long been recommended in discussions of PCa screening^5,23^, and PCPs are tasked with determining an individual’s risk based on factors such as his family history and ethnicity. However, physicians demonstrate different attitudes towards screening, with some screening all men proactively to avoid underdiagnoses, some screening only those men who request it, and some who attempt to weigh the costs and benefits of PSA screening on a case-by-case basis^24,25^. PCPs, who are already limited by time constraints and their patients’ other health issues, must carefully discuss the complex risks and benefits of PSA screening with their patients^26^.

Quantitative risk stratification could guide physicians in their screening conversations with patients by providing an objective risk-equivalent age for the development of higher-grade or other aggressive disease. This allows for simpler and more standardized informed decision-making regarding whether an individual man might benefit from PCa screening. For example, physicians who normally initiate discussions at some age (e.g., 50-55) could shift the timing of the PSA screening discussion according to the PCa risk-equivalent age. Some men might need to begin PCa screening at a relatively young age to detect an early-onset clinically-significant cancer. The PHS has previously demonstrated high PPV of PSA testing for aggressive PCa in men with progressively higher scores^9^. The potential utility of PCa risk-equivalent age in the clinic is additionally demonstrated by its impact on PPV. Suppose a 60-year old man presents to his physician to inquire about PCa screening. If this man has a PCa risk-equivalent age that is close to his true age (57-63), the PPV of a PSA test for him is approximately 30%; this is close to the overall PPV of PSA testing^5^. If his risk-equivalent age is <57, the PPV decreases to 10%, and he might be reassured in foregoing PSA testing. Postponing—or even forgoing—screening in men with low PHS percentiles to when they reach their risk-equivalent age could decrease the harms associated with screening, or early detection and treatment of PCa^4,5^. Other men may choose to delay the initiation of PSA testing until they are older and have increased risk. On the other hand, if this same man has a risk-equivalent age >63, the PPV of PSA testing increases substantially to 48%, implying that screening may be more informative for him. Identifying men with higher-grade disease at an early stage allows for definitive treatment to prevent cancer progression or metastases^11^.

Cost-effectiveness is another concern regarding PCa screening. Use of PHS, a one-time test valid for a man’s entire life, can improve screening efficiency while reducing overall costs. The genotyping chip assay requires only a saliva sample and can be run for costs similar to those for single-gene testing (e.g. the *BRCA* mutation). Genotyping also informs genetic risks for other diseases, possibly allowing multiple tests to be run on the same genotype results^27,28^. PSA screening (and subsequent prostate biopsy) could be offered only to those men at higher risk of aggressive PCa. PHS could increase the efficiency of any PCa screening program by incorporating knowledge that there are some men with higher baseline genetic risks of developing higher-grade PCa, even at a relatively young age, while others have a low baseline genetic risk.

Limitations of this work include that the PHS did not incorporate genotypic data from men of non-European ancestry during its development^9^, a reflection of the available data, which may affect the use of the PHS for screening decision-making in men from other ethnic groups. This is noteworthy, as disparities in PCa incidence and survival show that in the USA, men with African ancestry are more likely to develop PCa and to die from their disease^29^. Our group and others are studying the application of genetic scores to non-European ethnic groups. Additionally, there are now over 140 SNPs reported to have associations with PCa, identified using a meta-analysis that included ProtecT data^30^. However, not all of these SNPs are represented on the custom array used to develop the original PHS. Furthermore, the PHS model was validated using independent data from the ProtecT study; the inclusion of those other SNPs associated with PCa would have introduced circularity into the validation. Finally, we only evaluated higher-grade Gleason scores. For a more comprehensive assessment of clinically-significant PCa, future work should include other prognostically important clinical factors (including TNM stage, risk group stratification, and PSA at diagnosis).

We conclude that clinically-meaningful risk stratification could be achieved through application of a PHS that predicts of age of onset of higher-grade PCa to UK population data. PHS can also be used to calculate estimates of risk-equivalent age for the development of higher-grade PCa for individual men. The PPV of PSA was higher for men with higher PHS-adjusted PCa-equivalent ages. The assessment of personalized genetic risk via PHS could assist patients and physicians, alike, with the important decision of whether, and when, to initiate PCa screening.

## Supporting information

Supplemental Figure and PRACTICAL info

## Conflicts of Interest

All authors declare no support from any organization for the submitted work except as follows: AMD report a research grant from the US Department of Defense. OAA reports research grants from KG Jebsen Stiftelsen, Research Council of Norway, and South East Norway Health Authority.

Authors declare no financial relationships with any organizations that might have an interest in the submitted work in the previous three years except as follows, with all of these relationships outside the present study:

TMS reports honoraria from WebMD, Inc. for educational content, as well as a past research grant from Varian Medical Systems. ASK reports advisory board memberships for Sanofi-Aventis, Dendreon, and Profound. AK reports paid work for Certara Quantitative Systems Pharmacology.

Authors declare no other relationships or activities that could appear to have influenced the submitted work except as follows:

OAA has a patent application # U.S. 20150356243 pending; AMD also applied for this patent application and assigned it to UC San Diego. AMD has additional disclosures outside the present work: founder, equity holder, and advisory board member for CorTechs Labs, Inc.; advisory board member of Human Longevity, Inc.; recipient of nonfinancial research support from General Electric Healthcare.

Additional acknowledgments for the PRACTICAL consortium and contributing studies are described in the Supplemental Material.

## Funding for this study

This study was funded in part by a grant from the United States Department of Defense (#W81XWH-13-1-0391), the Research Council of Norway (#223273), KG Jebsen Stiftelsen, and South East Norway Health Authority. RMM is supported by a Cancer Research UK Programme Grant, the Integrative Cancer Epidemiology Programme (C18281/A19169), and the National Institute for Health Research (NIHR) Bristol Biomedical Research Centre. The views expressed are those of the author(s) and not necessarily those of the NIHR or the Department of Health and Social Care.

Funding for the PRACTICAL consortium member studies is detailed in the Supplemental Material.

A preliminary version of this work was presented in abstract form at the Genitourinary Cancers Symposium, February 14-16, 2019, San Francisco, CA, USA.

