## Supplemental Figure and PRACTICAL info for "A genetic hazard score to personalize prostate cancer screening, applied to population data"

**Supplemental eFigure 1**. Survival free of *any* prostate cancer, as derived from application of polygenic hazard score (PHS) hazard ratios and population data from the United Kingdom. The overall population incidence is taken as the median risk (50^th^ percentile). Hazard ratios were calculated within ProtecT data for various levels of genetic risk (i.e., PHS percentiles: 1, 5, 20, 80, 95, and 99) and used to adjust the median risk curve. Blue lines represent genetic risk lower than the median while red lines represent genetic risk higher than the median.

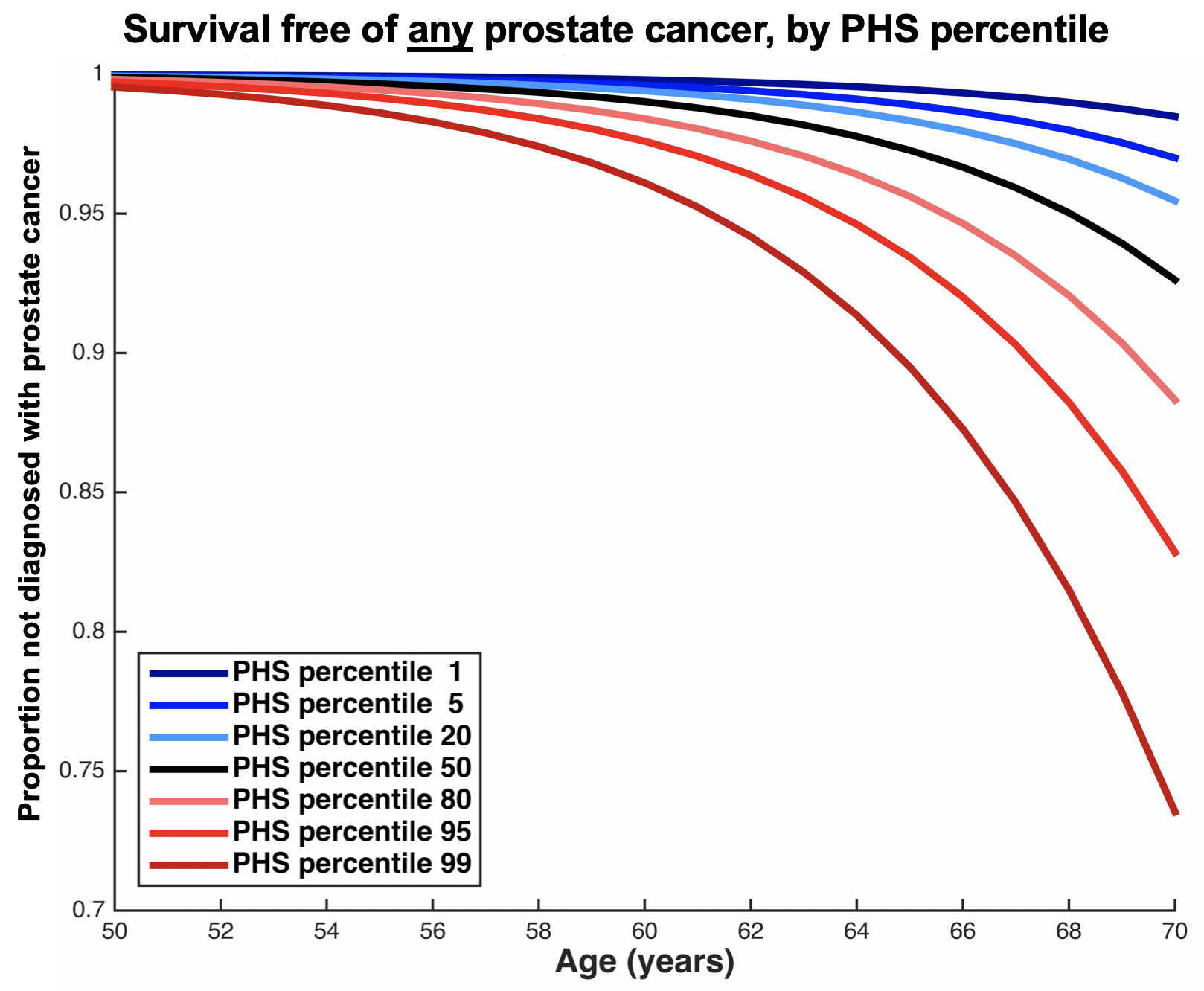

**Supplemental Material**

**The PRACTICAL CONSORTIUM (in addition to those named in the author list)**

Information of the consortium can be found at <http://practical.icr.ac.uk/>

Additional members from the consortium are:

Margaret Cook ^1^
Michelle Guy ^2^
Koveela Govindasami ^2^

Daniel Leongamornlert ^3^

Emma J. Sawyer ^2^

Rosemary Wilkinson ^2^

Edward J. Saunders ^2^

Malgorzata Tymrakiewicz ^2^

Tokhir Dadaev ^2^

Angela Morgan ^2^

Cyril Fisher ^2^

Steve Hazell ^2^

Naomi Livni ^2^

Artitaya Lophatananon ^4,5^

Robert Szulkin ^6,7^

Jan Adolfsson ^8,9^

Paer Stattin ^10,11^

Jan-Erik Johansson ^12^

Carin Cavalli-Bjoerkman ^13^

Ami Karlsson ^13^

Michael Broms ^13^

Anssi Auvinen ^14^

Paula Kujala ^15^

Kirsi Talala ^16^

Teemu Murtola ^17,18^

Kimmo Taari ^19^

Maren Weischer ^20^

Sune F. Nielsen ^20,21^

Peter Klarskov ^22^

Martin Andreas Røder ^23^

Peter Iversen ^23^

Hans Wallinder ^24^

Sven Gustafsson ^24^

Angela Cox ^25^

Paul Brown ^26^

Anne George ^27^

Gemma Marsden ^28^

Michael Davis ^29^

Wei Zheng ^30^

Lisa B. Signorello ^31^

William J. Blot ^32,33^

Lori Tillmans ^34^

Shaun Riska ^35^

Liang Wang ^36^

Antje Rinckleb ^37^

Jan Lubinski ^38^

Christa Stegmaier ^39^

Julio Pow-Sang ^40^

Hyun Park ^41^

Selina Radlein ^41^

Maria Rincon ^41^

James Haley ^41^

Babu Zachariah ^41^

Darina Kachakova ^42^

Elenko Popov ^43^

Atanaska Mitkova ^42^

Aleksandrina Vlahova ^44^

Tihomir Dikov ^44^

Svetlana Christova ^44^

Peter Heathcote ^45^

Glen Wood ^45^

Greg Malone ^45^

Pamela Saunders ^45^

Allison Eckert ^45^

Trina Yeadon ^45^

Kris Kerr ^45^

Angus Collins ^45^

Megan Turner ^45^

Srilakshmi Srinivasan ^45,46^

Mary-Anne Kedda ^45^

Kimberly Alexander ^45^

Tracy Omara ^45^

Huihai Wu ^47^

Rui Henrique ^48^

Pedro Pinto ^48^

Joana Santos ^48^

Joao Barros-Silva ^48^

Mohamed El Tibi ^49^

Graham G. Giles ^50,51^

Melissa C. Southey ^52^

Liesel M. Fitzgerald ^50,53^

John Pedersen ^54^

John L. Hopper ^51^

Robert MacInnis ^50,51^

Brian E. Henderson* ^55^

Fredrick Schumacher ^56,57^

Christopher A. Haiman ^58^

Janet L. Stanford ^59,60^

Susanne Kolb ^59^

Yong-Jie Lu ^61^

Hong-Wei Zhang ^62^

^1^ Centre for Cancer Genetic Epidemiology, Department of Public Health and Primary Care, University of Cambridge, Strangeways Research Laboratory, Cambridge CB1 8RN, UK

^2^ The Institute of Cancer Research, London, SM2 5NG, UK

^3^ Cancer Genome Project, Wellcome Trust Sanger Institute, Hinxton, CB10 1SA, UK

^4^ Institute of Population Health, University of Manchester, Manchester, UK

^5^ Warwick Medical School, University of Warwick, Coventry, UK

^6^ Division of Family Medicine, Department of Neurobiology, Care Science and Society, Karolinska Institutet, Huddinge, SE-171 77 Stockholm, Sweden

^7^ Scandinavian Development Services, Danderyd, 182 33, Sweden

^8^ Department of Clinical Science, Intervention and Technology, Karolinska Institutet, Stockholm, Sweden

^9^ Swedish Agency for Health Technology Assessment and Assessment of Social Services, Stockholm, Sweden

^10^ Department of Surgical and Perioperative Sciences, Urology and Andrology, Umea University, Umeå, Sweden

^11^ Department of Surgical Sciences, Uppsala University, Uppsala, Sweden

^12^ Department of Urology, Faculty of Medicine and Health, Örebro University, Örebro, Sweden

^13^ Department of Medical Epidemiology and Biostatistics, Karolinska Institute, Stockholm, Sweden

^14^ Department of Epidemiology, School of Health Sciences, FI-3314, University of Tampere, Tampere, Finland

^15^ Fimlab Laboratories, Tampere University Hospital, Tampere, Finland

^16^ Finnish Cancer Registry, Helsinki, Finland

^17^ Faculty of Medicine and Life Sciences, University of Tampere, Tampere, Finland

^18^ Department of Urology, Tampere University Hospital, Tampere, Finland

^19^ Department of Urology, Helsinki University Central Hospital and University of Helsinki, Helsinki, Finland

^20^ Department of Clinical Biochemistry, Herlev and Gentofte Hospital, Copenhagen University Hospital, Herlev, 220 Copenhagen, Denmark

^21^ Faculty of Health and Medical Sciences, University of Copenhagen, 2200 Copenhagen, Denmark

^22^ Department of Urology, Herlev and Gentofte Hospital, Copenhagen University Hospital, Herlev Ringvej 75, DK-2730 Herlev, Denmark

^23^ Copenhagen Prostate Cancer Center, Department of Urology, Rigshospitalet, Copenhagen University Hospital, DK-2730 Herlev, Copenhagen, Denmark

^24^ Department of Epidemiology and Biostatistics, School of Public Health, Imperial College, London, UK

^25^ Sheffield Institute for Nucleic Acids, University of Sheffield, Sheffield, UK

^26^ University of Cambridge, Department of Oncology, Box 279, Addenbrooke's Hospital, Hills Road Cambridge CB2 0QQ, UK

^27^ Cambridge Cancer Trials Centre, Cambridge Clinical Trials Unit - Cancer Theme, Cambridge University Hospitals NHS Foundation Trust, Box 279 (S4), Addenbrookes Hospital, Cambridge Biomedical Campus, Hills Road, Cambridge, CB2 0QQ, UK

^28^ Nuffield Department of Surgical Sciences, University of Oxford, Oxford, UK, Faculty of Medical Science, University of Oxford, John Radcliffe Hospital, Oxford, UK

^29^ School of Social and Community Medicine, Univerity of Bristol, Canynge Hall, 39 Whatley Road, Bristol BS8 2PS, UK

^30^ Division of Epidemiology, Department of Medicine, Vanderbilt University Medical Center, 2525 West End Avenue, Suite 800, Nashville, TN 37232 USA.

^31^ National Cancer Institute, NIH, 9609 Medical Center Drive, Suite 2W-172, MSC 9712, Bethesda, MD, 20892-9712 (mail), Rockville, MD 20850 (FedEx/Courier), USA

^32^ Division of Epidemiology, Department of Medicine, Vanderbilt University Medical Center, 2525 West End Avenue, Suite 600, Nashville, TN 37232 USA.

^33^ International Epidemiology Institute, Rockville, MD 20850, USA

^34^ Department of Laboratory Medicine and Pathology, Mayo Clinic, Rochester, MN 55905, USA.

^35^ Mayo Clinic, Rochester, Minnesota, USA

^36^ Taipei Medical University-Shuang-Ho Hospital

^37^ Department of Urology, University Hospital Ulm, Germany

^38^ International Hereditary Cancer Center, Department of Genetics and Pathology, Pomeranian Medical University, Szczecin, Poland

^39^ Saarland Cancer Registry, 66119 Saarbrücken, Germany

^40^ Genitourinary Program, Moffitt Cancer Center, 12902 Magnolia Drive, Tampa, FL 33612, USA

^41^ Department of Cancer Epidemiology, Moffitt Cancer Center, 12902 Magnolia Drive, Tampa, FL 33612, USA

^42^ Molecular Medicine Center, Department of Medical Chemistry and Biochemistry, Medical University, Sofia, 2 Zdrave Str., 1431 Sofia, Bulgaria

^43^ Department of Urology and Alexandrovska University Hospital, Medical University, Sofia, Bulgaria

^44^ Department of General and Clinical Pathology and Alexandrovska University Hospital, Medical University, Sofia, Bulgaria

^45^ Australian Prostate Cancer Research Centre-Qld, Institute of Health and Biomedical Innovation and School of Biomedical Science, Queensland University of Technology, Brisbane, Australia

^46^ Translational Research Institute, Brisbane, Queensland, Australia

^47^ The University of Surrey, Guildford, Surrey, GU2 7XH

^48^ Department of Genetics, Portuguese Oncology Institute, Porto, Portugal

^49^ University Hospital "Tsaritsa Yoanna", Medical University, Sofia, Bulgaria

^50^ Cancer Epidemiology & Intelligence Division, Cancer Council Victoria, 615 St Kilda Road, Melbourne, Victoria, 3004, Australia

^51^ Centre for Epidemiology and Biostatistics, Melbourne School of Population and Global Health, The University of Melbourne, Melbourne, Victoria 3010, Australia

^52^ Precision Medicine, School and Clinical Sciences at Monash Health, Monash University, Clayton, Victoria, 3168.

^53^ Cancer, Genetics and Immunology, Menzies Institute of Medical Research, Tasmania, 7000, Australia

^54^ Tissupath Pty Ltd., Melbourne,Victoria 3122, Australia

^55^ Department of Preventive Medicine, Keck School of Medicine, University of Southern California/Norris Comprehensive Cancer Center, Los Angeles, California, US

^56^ Department of Population and Quantitative Health Sciences, Case Western Reserve University, Cleveland, OH 44106-7219, USA

^57^ Seidman Cancer Center, University Hospitals, Cleveland, OH 44106, USA.

^58^ Department of Preventive Medicine, Keck School of Medicine, University of Southern California/Norris Comprehensive Cancer Center, Los Angeles, CA 90015, USA

^59^ Division of Public Health Sciences, Fred Hutchinson Cancer Research Center, Seattle, Washington, 98109-1024, USA

^60^ Department of Epidemiology, School of Public Health, University of Washington, Seattle, Washington 98195, USA

^61^ Centre for Molecular Oncology, Barts Cancer Institute, Queen Mary University of London, John Vane Science Centre, Charterhouse Square, London, EC1M 6BQ, UK

^62^ Second Military Medical University, 800 Xiangyin Rd., Shanghai 200433, P. R. China

* In memorium

**Funding for the CRUK study and PRACTICAL consortium:**

This work was supported by the Canadian Institutes of Health Research, European Commission's Seventh Framework Programme grant agreement n° 223175 (HEALTH-F2-2009-223175), Cancer Research UK Grants C5047/A7357, C1287/A10118, C5047/A3354, C5047/A10692, C16913/A6135, and The National Institute of Health (NIH) Cancer Post-Cancer GWAS initiative grant: No. 1 U19 CA 148537-01 (the GAME-ON initiative).

**COGS acknowledgement:**

This study would not have been possible without the contributions of the following: Per Hall (COGS); Douglas F. Easton, Paul Pharoah, Kyriaki Michailidou, Manjeet K. Bolla, Qin Wang (BCAC), Andrew Berchuck (OCAC), Rosalind A. Eeles, Douglas F. Easton, Ali Amin Al Olama, Zsofia Kote-Jarai, Sara Benlloch (PRACTICAL), Georgia Chenevix-Trench, Antonis Antoniou, Lesley McGuffog, Fergus Couch and Ken Offit (CIMBA), Joe Dennis, Alison M. Dunning, Andrew Lee, and Ed Dicks, Craig Luccarini and the staff of the Centre for Genetic Epidemiology Laboratory, Javier Benitez, Anna Gonzalez-Neira and the staff of the CNIO genotyping unit, Jacques Simard and Daniel C. Tessier, Francois Bacot, Daniel Vincent, Sylvie LaBoissière and Frederic Robidoux and the staff of the McGill University and Génome Québec Innovation Centre, Stig E. Bojesen, Sune F. Nielsen, Borge G. Nordestgaard, and the staff of the Copenhagen DNA laboratory, and Julie M. Cunningham, Sharon A. Windebank, Christopher A. Hilker, Jeffrey Meyer and the staff of Mayo Clinic Genotyping Core Facility

Funding for the iCOGS infrastructure came from: the European Community's Seventh Framework Programme under grant agreement n° 223175 (HEALTH-F2-2009-223175) (COGS), Cancer Research UK (C1287/A10118, C1287/A 10710, C12292/A11174, C1281/A12014, C5047/A8384, C5047/A15007, C5047/A10692, C8197/A16565), the National Institutes of Health (CA128978) and Post-Cancer GWAS initiative (1U19 CA148537, 1U19 CA148065 and 1U19 CA148112 - the GAME-ON initiative), the Department of Defence (W81XWH-10-1-0341), the Canadian Institutes of Health Research (CIHR) for the CIHR Team in Familial Risks of Breast Cancer, Komen Foundation for the Cure, the Breast Cancer Research Foundation, and the Ovarian Cancer Research Fund.

**Additional funding and acknowledgments from studies in PRACTICAL:**

The Department of Medical Epidemiology and Biostatistics, Karolinska Institute, Stockholm, Sweden was supported by the Cancer Risk Prediction Center (CRisP; www.crispcenter.org), a Linneus Centre (Contract ID 70867902) financed by the Swedish Research Council, Swedish Research Council (grant no K2010-70X-20430-04-3), the Swedish Cancer Foundation (grant no 09-0677), the Hedlund Foundation, the Soederberg Foundation, the Enqvist Foundation, ALF funds from the Stockholm County Council. Stiftelsen Johanna Hagstrand och Sigfrid Linner's Minne, Karlsson's Fund for urological and surgical research. We thank and acknowledge all of the participants in the Stockholm-1 study. We thank Carin Cavalli-Bjoerkman and Ami Roennberg Karlsson for their dedicated work in the collection of data. Michael Broms is acknowledged for his skilful work with the databases. KI Biobank is acknowledged for handling the samples and for DNA extraction. Hans Wallinder at Aleris Medilab and Sven Gustafsson at Karolinska University Laboratory are thanked for their good cooperation in providing historical laboratory results.

The coordination of EPIC was financially supported by the European Commission (DG-SANCO) and the International Agency for Research on Cancer. The national cohorts (that recruited male participants) are supported by Danish Cancer Society (Denmark); German Cancer Aid, German Cancer Research Center (DKFZ), Federal Ministry of Education and Research (BMBF), Deutsche Krebshilfe, Deutsches Krebsforschungszentrum and Federal Ministry of Education and Research (Germany); the Hellenic Health Foundation (Greece); Associazione Italiana per la Ricerca sul Cancro-AIRC-Italy and National Research Council (Italy); Dutch Ministry of Public Health, Welfare and Sports (VWS), Netherlands Cancer Registry (NKR), LK Research Funds, Dutch Prevention Funds, Dutch ZON (Zorg Onderzoek Nederland), World Cancer Research Fund (WCRF), Statistics Netherlands (The Netherlands); Health Research Fund (FIS), PI13/00061 to Granada; , PI13/01162 to EPIC-Murcia), Regional Governments of Andalucía, Asturias, Basque Country, Murcia and Navarra, ISCIII RETIC (RD06/0020) (Spain); Swedish Cancer Society, Swedish Research Council and County Councils of Skåne and Västerbotten (Sweden); Cancer Research UK (14136 to EPIC-Norfolk;

C570/A16491 and C8221/A19170 to EPIC-Oxford), Medical Research Council

(1000143 to EPIC-Norfolk, MR/M012190/1 to EPIC-Oxford) (United Kingdom).

The ESTHER study was supported by a grant from the Baden Württemberg Ministry of Science, Research and Arts. Additional cases were recruited in the context of the VERDI study, which was supported by a grant from the German Cancer Aid (Deutsche Krebshilfe). The ESTHER group would like to thank Hartwig Ziegler, Sonja Wolf, Volker Hermann, Katja Butterbach for valuable contributions to the study.

The FHCRC studies were supported by grants RO1CA056678, RO1CA082664, and RO1CA092579 from the US National Cancer Institute, National Institutes of Health, with additional support from the Fred Hutchinson Cancer Research Center. We thank all the men that participated in these studies.

The IPO-Porto study was funded by Fundação para a Ciência e a Tecnologia (FCT; UID/DTP/00776/2013 and PTDC/DTP-PIC/1308/2014) and by IPO-Porto Research Center (CI-IPOP-16-2012 and CI-IPOP-24-2015). MC and MPS are research fellows from Liga Portuguesa Contra o Cancro, Núcleo Regional do Norte. SM is a research fellow from FCT (SFRH/BD/71397/2010). We would like to express our gratitude to all patients and families who have participated in this study.

The Mayo group was supported by the US National Cancer Institute (R01CA72818)

The Melbourne Collaborative Cohort Study (MCCS) cohort recruitment was funded by VicHealth and Cancer Council Victoria. The MCCS was further supported by Australian National Health and Medical Research Council grants 209057 and 396414 and by infrastructure provided by Cancer Council Victoria. Cases and their vital status were ascertained through the Victorian Cancer Registry and the Australian Institute of Health and Welfare, including the National Death Index and the Australian Cancer Database.

The MEC was supported by NIH grants CA63464, CA54281, CA098758, and CA164973.

The Moffitt group was supported by the US National Cancer Institute (R01CA128813, PI: J.Y. Park).

The PCMUS study was supported by the Bulgarian National Science Fund, Ministry of Education and Science (contract DOO-119/2009; DUNK01/2-2009; DFNI-B01/28/2012) with additional support from the Science Fund of Medical University - Sofia (contract 51/2009; 8I/2009; 28/2010).

ProtecT would like to acknowledge the support of The University of Cambridge, Cancer Research UK. Cancer Research UK grants [C8197/A10123] and [C8197/A10865] supported the genotyping team. We would also like to acknowledge the support of the National Institute for Health Research which funds the Cambridge Bio-medical Research Centre, Cambridge, UK. We would also like to acknowledge the support of the National Cancer Research Prostate Cancer: Mechanisms of Progression and Treatment (PROMPT) collaborative (grant code G0500966/75466) which has funded tissue and urine collections in Cambridge. We are grateful to staff at the Welcome Trust Clinical Research Facility, Addenbrooke’s Clinical Research Centre, Cambridge, UK for their help in conducting the ProtecT study. We also acknowledge the support of the NIHR Cambridge Biomedical Research Centre, the DOH HTA (ProtecT grant) and the NCRI / MRC (ProMPT grant) for help with the bio-repository. The UK Department of Health funded the ProtecT study through the NIHR Health Technology Assessment Programme (projects 96/20/06, 96/20/99). The ProtecT trial and its linked ProMPT and CAP (Comparison Arm for ProtecT) studies are supported by Department of Health, England; Cancer Research UK grant number C522/A8649, Medical Research Council of England grant number G0500966, ID 75466 and The NCRI, UK. The epidemiological data for ProtecT were generated though funding from the Southwest National Health Service Research and Development. DNA extraction in ProtecT was supported by USA Dept of Defense award W81XWH-04-1-0280, Yorkshire Cancer Research and Cancer Research UK. The authors would like to acknowledge the contribution of all members of the ProtecT study research group. The views and opinions expressed therein are those of the authors and do not necessarily reflect those of the Department of Health of England. The bio-repository from ProtecT is supported by the NCRI (ProMPT) Prostate Cancer Collaborative and the Cambridge BMRC grant from NIHR. We acknowledge support from the National Cancer Research Institute (National Institute of Health Research (NIHR) Collaborative Study: “Prostate Cancer: Mechanisms of Progression and Treatment (PROMPT)” (grant G0500966/75466). We thank the National Institute for Health Research, Hutchison Whampoa Limited, the Human Research Tissue Bank (Addenbrooke’s Hospital), and Cancer Research UK. The authors would like to thank those men with prostate cancer and the subjects who have donated their time and their samples to the Cambridge Biorepository, which were used in this research. We also would like to acknowledge to support of the research staff in S4 who so carefully curated the samples and the follow-up data (Jo Burge, Marie Corcoran, Anne George, and Sara Stearn).

The CAP trial is funded by Cancer Research UK and the UK Department of Health (C11043/A4286, C18281/A8145, C18281/A11326, and C18281/A15064), with the University of Bristol as sponsor.  The ProtecT trial is funded by the UK National Institute for Health Research (NIHR), Health Technology Assessment Programme (projects 96/20/06, 96/20/99), with the University of Oxford as sponsor. <http://www.nets.nihr.ac.uk/projects/hta/962099>).

The QLD research is supported by The National Health and Medical Research Council (NHMRC) Australia Project Grants [390130, 1009458] and NHMRC Career Development Fellowship, Cancer Australia PdCCRS and Cancer Council Queensland funding to J Batra. The QLD team would like to acknowledge and sincerely thank the urologists, pathologists, data managers and patient participants who have generously and altruistically supported the QLD cohort.

The Australian Prostate Cancer BioResource (APCB) was supported by The National Health and Medical Research Council, Enabling Grant [614296] and the Prostate Cancer Foundation of Australia.The Australian Prostate Cancer BioResource (APCB) would like to acknowledge and sincerely thank the urologists, pathologists, coordinators, data managers, nurses and patient participants who have generously and altruistically supported the APCB.

SCCS is funded by NIH grant R01 CA092447, and SCCS sample preparation was conducted at the Epidemiology Biospecimen Core Lab that is supported in part by the Vanderbilt-Ingram Cancer Center (P30 CA68485). Data on SCCS cancer cases used in this publication were provided by the Alabama Statewide Cancer Registry; Kentucky Cancer Registry, Lexington, KY; Tennessee Department of Health, Office of Cancer Surveillance; Florida Cancer Data System; North Carolina Central Cancer Registry, North Carolina Division of Public Health; Georgia Comprehensive Cancer Registry; Louisiana Tumor Registry; Mississippi Cancer Registry; South Carolina Central Cancer Registry; Virginia Department of Health, Virginia Cancer Registry; Arkansas Department of Health, Cancer Registry, 4815 W. Markham, Little Rock, AR 72205. The Arkansas Central Cancer Registry is fully funded by a grant from National Program of Cancer Registries, Centers for Disease Control and Prevention (CDC). Data on SCCS cancer cases from Mississippi were collected by the Mississippi Cancer Registry which participates in the National Program of Cancer Registries (NPCR) of the Centers for Disease Control and Prevention (CDC). The contents of this publication are solely the responsibility of the authors and do not necessarily represent the official views of the CDC or the Mississippi Cancer Registry.

SEARCH is funded by a programme grant from Cancer Research UK [C490/A10124] and supported by the UK National Institute for Health Research Biomedical Research Centre at the University of Cambridge. The University of Cambridge has received salary support in respect of PP from the NHS in the East of England through the Clinical Academic Reserve.

The Tampere (Finland) study was supported by the Academy of Finland (251074), The Finnish Cancer Organisations, Sigrid Juselius Foundation, and the Competitive Research Funding of the Tampere University Hospital (X51003). The PSA screening samples were collected by the Finnish part of ERSPC (European Study of Screening for Prostate Cancer). TAMPERE would like to thank Riina Liikanen, Liisa Maeaettaenen and Kirsi Talala for their work on samples and databases.

UKGPCS would also like to thank the following for funding support: The Institute of Cancer Research and The Everyman Campaign, The Prostate Cancer Research Foundation, Prostate Research Campaign UK (now Prostate Action), The Orchid Cancer Appeal, The National Cancer Research Network UK, The National Cancer Research Institute (NCRI) UK. We are grateful for support of NIHR funding to the NIHR Biomedical Research Centre at The Institute of Cancer Research and The Royal Marsden NHS Foundation Trust. UKGPCS should also like to acknowledge the NCRN nurses, data managers and Consultants for their work in the UKGPCS study. UKGPCS would like to thank all urologists and other persons involved in the planning, coordination, and data collection of the study. Kenneth Muir as part of the UKGPCS study was supported by a CRUK programme grant, the Integrative Cancer Epidemiology Programme (C18281/A19169) and by the NIHR Manchester Biomedical Research Centre.

The Ulm group received funds from the German Cancer Aid (Deutsche Krebshilfe).

The Keith and Susan Warshaw Fund, C. S. Watkins Urologic Cancer Fund and The Tennity Family Fund supported the Utah study. The project was supported by Award Number P30CA042014 from the National Cancer Institute.
